# *In situ* visualization of autophagy suggests vesicle fusion can contribute to phagophore expansion

**DOI:** 10.64898/2026.03.29.715079

**Authors:** Claire Ortmann de Percin Northumberland, Mariya Licheva, Rahel Dabrowski, Rubén Gómez-Sánchez, Sabrina Berkamp, Philipp Schönnenbeck, Martin Graef, Claudine Kraft, Carsten Sachse

## Abstract

The autophagy core machinery mediates the enclosure of cytosolic cargo destined for degradation in the lysosome. The Atg9-Atg2-Atg18 complex coordinates phagophore expansion via directed lipid transfer until closure of the phagophore rim. Using an Atg2 variant (Atg2-PM4) as a model of decelerated autophagosome biogenesis, we visualized the morphological states prior to autophagosome closure by cryogenic correlative light and electron microscopy in *S. cerevisiae*. Using *in situ* cryo-electron tomography, we find an enlarged rim morphology of an expanding phagophore in Atg2-PM4 cells in comparison with Atg2 wildtype condition. Analysis of segmented rim membrane features as well as surrounding and attached vesicles suggest that the enlarged rims are a result of cytosolic vesicles fusing with the growing phagophore. High-resolution imaging in this study shows that, apart from the initial nucleation phase, vesicle fusion can also contribute to phagophore expansion during later stages of autophagosome biogenesis.

## Introduction

Autophagy is a conserved homeostatic process that recycles molecular components of cellular material such as damaged organelles and protein aggregates by delivering them to the yeast and plant vacuole or the mammalian lysosome (Takeshige et al., 1992; Wen and Klionsky, 2016). The autophagic process can be triggered in response to cellular stresses such as nutrient deprivation (Takeshige et al., 1992). There are three main types of autophagy: macroautophagy, microautophagy, and chaperone-mediated autophagy (Xie and Klionsky, 2007). The morphological hallmark of macroautophagy (hereafter autophagy) is the *de novo* formation of a double-membrane compartment, the autophagosome, enclosing the autophagic cargo (Wen and Klionsky, 2016). By now, over 40 autophagy-related (Atg) proteins have been identified, initially in *Saccharomyces (S.) cerevisiae* (Tsukada and Ohsumi, 1993) while approximately 18 belong to the core autophagy machinery (Kawabata and Yoshimori, 2020; Nakatogawa, 2020). The autophagy machinery is organized in the following six evolutionarily conserved protein complexes: Atg1 complex (Atg1, Atg13, Atg17, Atg29, Atg31), phosphatidylinositol (PI)-3 kinase complex (Atg6, Atg14, Vps34, Vps15, Atg38), Atg9, Atg2-Atg18 complex, Atg12-Atg5 conjugation system (Atg12, Atg7, Atg10, Atg5, Atg16), and Atg8 conjugation system (Atg8, Atg7, Atg3, Atg4) (Suzuki and Ohsumi, 2010; Noda, 2023; Nakatogawa, 2020). Autophagosome biogenesis temporally proceeds in five stages: phagophore initiation, nucleation, expansion, closure into an autophagosome, and autophagosome fusion with the vacuole or lysosome (Wen and Klionsky, 2016).

After initiation in *S. cerevisiae*, the Atg1 complex forms the phagophore assembly site (PAS) at the surface of the vacuole to nucleate downstream assemblies of PI3-kinase and Atg9-Atg2-Atg18 complexes (Fujioka et al., 2020; Hollenstein et al., 2019). Atg9 is stored in Golgi-derived cytosolic vesicles prior to fusion at the PAS (Noda et al., 2000; Yamamoto et al., 2012). At the early stages of autophagy during nucleation, a subset of Atg9 vesicles localizes to the PAS and constitutes an initial membrane seed to form a phagophore (Sawa-Makarska et al., 2020; Olivas et al., 2023; Mari et al., 2010). COPII vesicles have also been implicated in the lipid supply to the phagophore (Graef et al., 2013; Shima et al., 2019). The growing phagophore (also called isolation membrane) is derived from a cup-shaped membrane cisterna (Bieber et al., 2022; Shibutani and Yoshimori, 2014). During intermediate stages of autophagosome biogenesis, the core autophagy proteins Atg2 and Atg9 contribute to phagophore expansion: Atg2 by establishing membrane contact sites between the ER/ERES and the growing phagophore (together with TRAPPIII complex) and mediating lipid transfer from a donor membrane (Gómez-Sánchez et al., 2025, 2018; Kotani et al., 2018), and Atg9 by facilitating their distribution across both membrane leaflets (Chumpen Ramirez et al., 2023; Maeda et al., 2019; Matoba et al., 2020; Valverde et al., 2019). Atg9 is the only integral transmembrane protein among the core autophagy machinery components (Matoba et al., 2020; Noda et al., 2000) and was found to trimerize. The scramblase activity of Atg9 essential for autophagy (Maeda et al., 2020; Matoba et al., 2020) is likely to contribute to maintaining previously reported lipid asymmetry in autophagic membranes, with higher phosphatidylinositol-3-phosphate (PI3P) levels in the luminal leaflets of both the outer and inner membranes than in the cytoplasmic leaflet of the outer membrane or the inner leaflet of the inner membrane (Cheng et al., 2014). Atg2 is an elongated rod-shaped β-helix protein found to supply lipids by bridging the phagophore to donor membranes, e.g., ER, and providing a hydrophobic slide along its long axis that allows lipids to transfer from the N to the C-terminal end of the rod (Chowdhury et al., 2018; Valverde et al., 2019). It was shown that Atg2 has no lipid head-group transfer specificity (Osawa et al., 2019). A lipid transfer rate of 200 lipids per Atg2 molecule per second has been estimated *in vivo* using fluorescence microscopy (FM) to contribute to phagophore expansion (Dabrowski et al., 2023). It was also found that Vps13, a homologue of Atg2 supports the lipid transfer to phagophores to render it non-rate limiting (Dabrowski et al., 2023).

While the biochemical activities of both Atg2 and Atg9 in the context of autophagy have been characterized, how their respective functions are spatially and temporally coordinated to achieve a unidirectional lipid transfer to the growing phagophore remains under investigation. Two hypotheses have been put forward; the first one involving the *S. cerevisiae* long chain fatty acyl-CoA synthetase 1 (Faa1) and the second is based on two human scramblases TMEM41B and VMP1. Faa1 was observed localizing to phagophores and generating acyl-CoA using ATP (Schütter et al., 2020). Subsequently, newly synthesized acyl-CoA is channeled to the phospholipid synthesizing enzymes of the ER for *de novo* lipid synthesis. Newly synthesized lipids are subsequently transferred to the phagophore by Atg2. It has been proposed that *de novo* lipid synthesis increases the chemical potential and thereby providing a driving force for the unidirectional lipid transfer from the ER into the phagophore (Schütter et al., 2020). The second hypothesis is based on the findings that deleting the two human scramblases TMEM41B and VMP1 resulted in depletion of autophagic flux (Morita et al., 2018) in addition to the described interaction with ATG2A at the ER (Ghanbarpour et al., 2021) providing a driver of unidirectional lipid transfer to the phagophore. In addition to the described mechanisms of lipid transfer, other cellular processes may contribute to phagophore growth.

Phagophore expansion is further accompanied by the conjugation of ubiquitin-like Atg8 to the phagophore membrane. The responsible machinery is composed of Atg5-Atg12/Atg16 (Romanov et al., 2012) and acts like an E3 ligase complex to transfer Atg8 from Atg3 to membrane-resident phosphatidylethanolamine (Hanada et al., 2007). Phagophore expansion through lipids and Atg8 conjugation proceeds until the phagophore rims are ready to be fused to form a closed autophagosome. Deletion of *S. cerevisiae SNF7* as well as human *CHMP2B* and *VPS4* led to an accumulation of unsealed autophagosomes and consistently impaired autophagosome completion (Takahashi et al., 2018; Zhou et al., 2019). Snf7 and CHMP2B belong to the family of ESCRT-III proteins enabling membrane fusion and/or fission while in this case they mediate the closure of phagophores into double-membrane autophagosomes.

The interaction between Atg2 and Atg9 has also been shown to be essential for autophagic turn-over (Gómez-Sánchez et al., 2018). When disrupting the Atg2-Atg9 interaction using point mutations at the C-terminal region of Atg2 (named Atg2-PM4), both Atg9 and Atg2 still concentrate at the phagophore membrane, but in different locations. While Atg9 and Atg2 both localize to the phagophore rims, mutant Atg2-PM4 mislocalizes, showing distribution along the phagophore membrane distal from the rims (Gómez-Sánchez et al., 2018). It was suggested that as a result the association of ER and phagophore is altered (Gómez-Sánchez et al., 2018). Here, we characterized this mutant by live-cell fluorescence microscopy revealing delayed phagophore formation. Moreover, we visualized the native accumulated phagophores in the Atg2-PM4 mutant in *S. cerevisiae* using *in situ* cryogenic electron tomography (cryo-ET) coupled with cryogenic correlative light and electron microscopy (cryo-CLEM) and observed significantly enlarged phagophore rims upon disruption of the Atg2-Atg9 interaction. The tomograms provide evidence of vesicular fusion with the phagophore and suggest that this process can contribute to lipid growth at intermediate stages of autophagosome biogenesis during phagophore expansion.

## Results

### Phagophore formation is decelerated in Atg2-PM4 expressing cells

To better understand phagophore expansion and morphology, we focused on the Atg9-Atg2 interaction and turned to the previously characterized Atg2-PM4 mutant (Gómez-Sánchez et al., 2018) that carries five mutations in the C-terminal region, i.e. QKFST to AAAAA (1264-1268) (highlighted in AlphaFold 3 model; **Fig. S1**), disrupting the interaction of Atg2 with Atg9 at the phagophore. Therefore, we used live-cell fluorescence imaging to characterize autophagosome biogenesis and imaged phagophores using mCherry-Atg8 as described previously (Dabrowski et al., 2023) together with Atg2-WT-3xGFP and Atg2-PM4-3xGFP, respectively (**Fig. 1a**). Interestingly, in Atg2-PM4 cells we were unable to detect Atg2-PM4 puncta as opposed to the Atg2-WT control, which is consistent with previous studies suggesting that Atg2-PM4 mutant distributes across the phagophore membrane rather than accumulates at the phagophore rims (Gómez-Sánchez et al., 2018). When counting the number of Atg8 puncta per cell, we observed a slight decrease for Atg2-PM4 cells while the overall variation is very similar to that of Atg2-WT cells (**Fig. 1b**, left). More significantly, on average we found less than one closed phagophores per five cells, i.e., 0.2 phagophores per cell, in Atg2-PM4 compared with 0.7 per cell in Atg2-WT (**Fig. 1b**, right). These results indicate that the dynamic rates of autophagosome biogenesis are reduced in the Atg2-PM4 mutant. To confirm this hypothesis, we performed timelapse fluorescence microscopy and followed Atg8 puncta from nucleation until autophagosome fusion with the vacuole, for a maximum of 30 minutes (as previously described in (Dabrowski et al., 2023)). We observed an increase in the total lifetime of Atg8 positive structures in Atg2-PM4 mutant cells when compared with Atg2-WT cells, on average lasting a duration of 22 vs. 10 min in the Atg2-WT (**Fig. 1c**). Taken together, the fluorescence analyses of the Atg2-PM4 mutant revealed that autophagosome biogenesis was decelerated resulting in lower numbers of autophagosomes than in the Atg2-WT.

**Figure 1.**
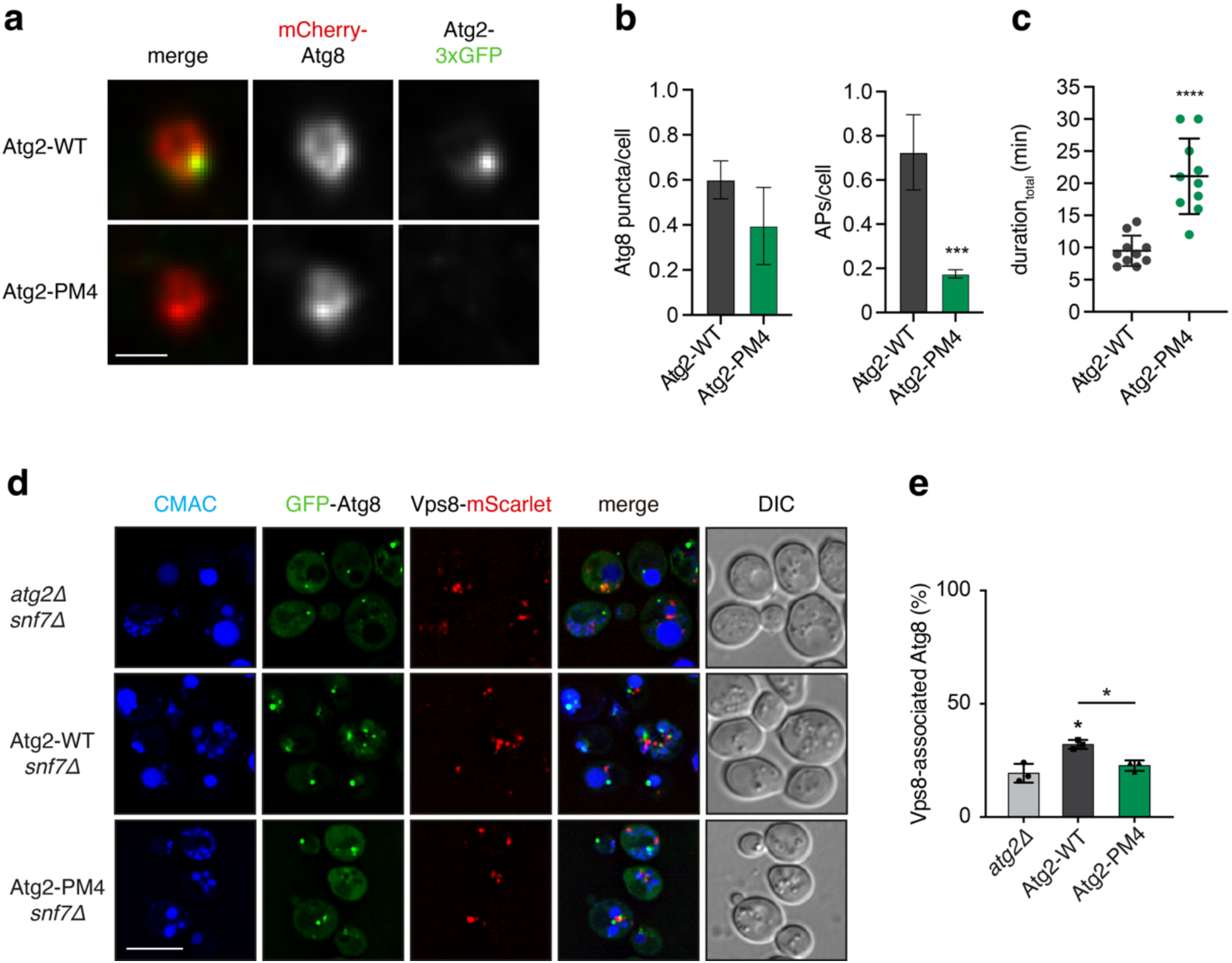
Fluorescence light microscopy live-cell imaging of autophagosome biogenesis in Atg2-PM4 cells and imaging of Atg8 and Vps8-positive structures. **a**) Representative live-cell fluorescence microscopy images of cells expressing Atg2 (Atg2-WT) or the Atg2-PM4 mutant along with mCherry-Atg8 (W303 strain). Images were acquired after 1 h of nitrogen starvation. **b**) Quantifications of data shown in panel a. The relative number of Atg8 puncta and autophagosomal/phagophore structures (APs) per cell of indicated strains was analyzed. (n=4; 200 structures analyzed in total/strain). T-test. **c**) Quantification of live-cell timelapse fluorescent images of cells expressing Atg2-WT or the Atg2-PM4 mutant along with mCherry-*ATG8*. Images were acquired after 1 h of nitrogen starvation. Duration_total_ describes the time from nucleation until fusion with the vacuole, as described previously (Dabrowski et al., 2023). (n=2; 10 events/strain). T-test. *P*-values for all Quantifications: *** *p*<0.0002; **** *p*>0.0001. **d**) The *snf7Δ atg2Δ* strain (BY) expressing GFP-Atg8 and Vps8-mScarlet (RGY1381,) was transformed with an empty vector (pRS416; *atg2Δ*) or plasmids expressing Atg2 or Atg2-PM4 mutant. Transformed cells were cultured in SD-URA to early log phase and transferred into SD-N starvation medium for 2 h to induce autophagy before being analyzed by live-cell imaging. **e**) Percentage of GFP-Atg8-labelled PAS associated with Vps8-mScarlet foci in the experiment shown in panel d). Black asterisk highlights significant differences between *atg2Δ* and Atg2 cells and grey asterisk indicates significant difference between Atg2 and Atg2-PM4 cells, significant meaning p < 0.05. Quantification in panel b) represents the average of three independent experiments ±SD. DIC, differential interference contrast. Scale bars: 1 µm (panel a) 4 µm (panel d).

### Establishing the *snf7Δ* yeast strain to accumulate phagophores

In order to enrich phagophore expansion events for down-stream cryo-electron microscopy studies, we turned to a *SNF7* deletion (*snf7Δ)* strain that was shown to accumulate open, i.e. unsealed phagophores at the vacuole (Zhou et al., 2019). At the same time, it had been characterized by vesicle trafficking defects and large endosomal structures named class E compartments (Raymond et al., 1992). We wanted to verify that these altered membrane dynamics did not affect phagophore biogenesis. To exclude this possibility, we therefore followed Vps8 foci to visualize class E compartments as it was shown to concentrate in these clusters upon ESCRT-III dysfunction (Russell et al., 2012). We turned to live-cell imaging to monitor Vps8-mScarlet foci alongside GFP-Atg8 in *atg2Δ*, Atg2-WT, and Atg2-PM4 cells with *snf7Δ* background, respectively (**Fig. 1d)**. When we estimated the fraction of Vps8 spatially close to Atg8, we observed approximately 25 % of Atg8-positive structures associated with class E compartments between the different strains (showing 20, 32 and 23 % association for *atg2Δ*, Atg2-WT, and Atg2-PM4 cells, respectively) (**Fig. 1e)**. As expected, we observed the presence of larger GFP-Atg8 foci in both Atg2-WT and Atg2-PM4 cells, when compared with *atg2Δ* cells, in line with previous studies (Gómez-Sánchez et al., 2018; Chumpen Ramirez et al., 2023). These differences in fluorescence intensity and clustering between Atg2-WT and Atg2-PM4 cells can be explained by the defective recruitment of Atg18 to the nascent phagophore in Atg2-PM4 cells (Gómez-Sánchez et al., 2018). Taken together, these live-cell imaging data indicate that the growing phagophores in the Atg2-PM4 mutant of the *snf7Δ* strain do not show increased association with class E compartments supporting the conclusion that they present a suitable experimental model to further visualize phagophore biogenesis.

### Visualization of phagophores in *SNF7* deletion cells by *in situ cryo-ET*

In order to study expanding phagophores of *S. cerevisiae* at higher resolution, we employed *in situ* cryo-electron tomography and continued to work with the *snf7Δ* strain (**Fig. 2a**). To identify the region of interest for tomographic data acquisition, we were guided by Atg9-3xmTagBFP2 positive puncta using cryo-CLEM. When we detected a BFP signal to validate the presence of Atg9, we generated cellular lamellae with a focused ion beam milling scanning electron microscope (FIB-SEM) operated at cryogenic temperatures (**Fig. 2b**) and correlated in-chamber fluorescence images after milling (**Fig. 2c**) with TEM overview images (**Fig. 2d-e**). In this way, we acquired a total of 13 tomograms of approximately 150-200 nm thickness and visualized six sections through phagophores in the presence of common autophagy factors including wildtype Atg2 (Atg2-WT) (**Fig. 2f, Table 1**). We observed characteristic phagophores of 300 – 400 nm in diameter with their closely apposed bilayers, in proximity to the ER and lipid droplets (**Fig. 2g-h**). Occasionally, extended densities were identified as membrane connections with approx. 20 to 24 nm in length (**Table 2**) bridging the phagophore membrane to putative donor membranes, i.e., ER (**Fig. 2g**\*) and lipid droplet surfaces (**Fig. 2h**\*). These densities match the length of the mammalian Atg2 and yeast Vps13p molecules (Valverde et al., 2019; De et al., 2017). The phagophores imaged in the Atg2-WT *snf7Δ* strain (**Fig. 2g-i**) are consistent with previously observed phagophore ultrastructures in *ypt7Δ* cells with the characteristic cup-shaped membrane as well as protein-mediated membrane contact sites with the phagophore (Bieber et al., 2022). As phagophores accumulate in the cytosol of *snf7Δ* cells and are prevented from fusing with the vacuole, we successfully observed multiple stages of phagophore expansion in Atg2-WT conditions of this otherwise highly transient process.

**Figure 2:**
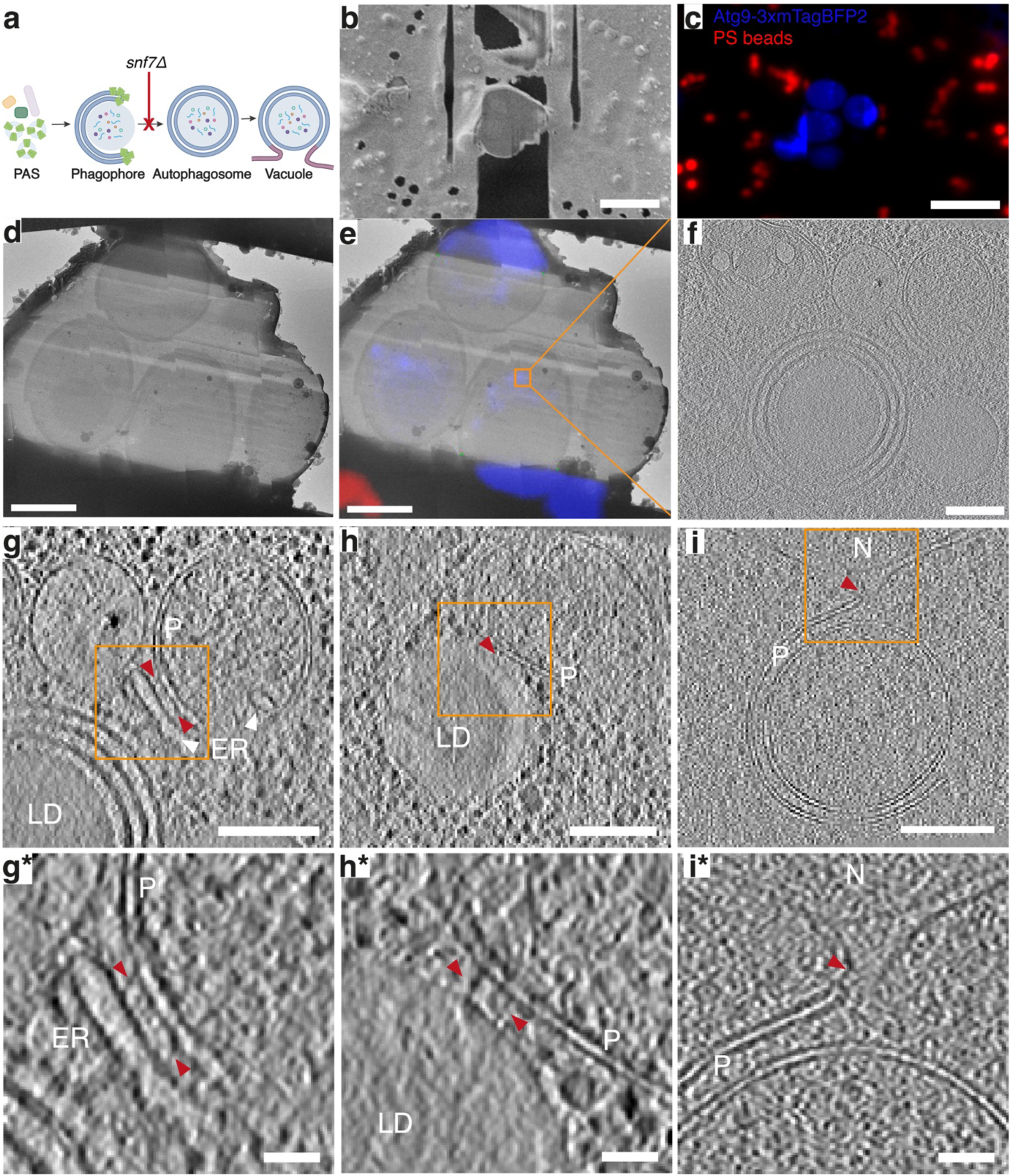
Visualization of yeast phagophores by correlative light and electron microscopy. **a**) Schematic representation of autophagosome biogenesis including phagophore closure mediated by Snf7 that was deleted for the analysis. **b**) SEM image of yeast cellular lamella used to characterize wild-type phagophore elongation in *snf7Δ* cells. **c**) Same lamella as in panel b imaged in a fluorescence microscope under cryogenic conditions. Signal shown in blue represents Atg9-3xmTagBFP2 and signal shown in red originates from polystyrene (PS) fiducial beads. **d**) TEM overview of the same lamella as shown in panels b and c. **e**) TEM overview of lamella (panel d) correlated with the fluorescence image of same lamella (panel c). Region on which tilt series was acquired is indicated by orange box. **f**) Tomographic slice of multiple tomograms showing phagophore, acquired from the region indicated in panel e. **g**)-**i**) Tomographic slices of phagophores from *snf7Δ* yeast cells, denoised with IsoNet (Liu et al., 2022). **g\***)-**i\***) represent enlarged views of regions from panels g) to i) marked with orange boxes. Red arrows in panels g) to i*) show densities spanning from a donor membrane to the phagophore membrane, measured lengths listed in Table 2. Phagophores are indicated with P, lipid droplets with LD, endoplasmic reticulum with ER, vacuole with V, nucleus with N. Scale bars b)-c) 10 µm, d)-e) 3 µm, f)-i) 200 nm, and g*)-i*) 100nm.

**Table 1:** Summary of cryo-electron tomography data collection.

|  | Total tomograms collected | Tomograms containing phagophore rims | Acceleration voltage (keV) | Cell culture batch |
| --- | --- | --- | --- | --- |
| Total | 30 | 17 |  |  |
| Atg2-WT | 13 | 2 | 200 | Batch 1 |
|  |  | 4 | 300 | Batch 2 |
| Atg2-PM4 | 17 | 4 | 200 | Batch 1 |
|  |  | 7 | 200 | Batch 2 |

**Table 2:** Length of densities tethering phagophore and putative donor membranes in both Atg2-WT and Atg2-PM4 cells.

| Phenotype | Measured length (nm) | Figure |
| --- | --- | --- |
| Atg2-WT | 20 | Fig. 2g* (top) |
|  | 21 | Fig. 2g* (bottom) |
|  | 24 | Fig. 2h* (top) |
|  | 23 | Fig. 2h* (bottom) |
| Atg2-PM4 | 29 | Fig. 3h* (top) |
|  | 29 | Fig. 3h* (bottom) |
|  | 23 | Fig. 3j |

### Phagophore morphology is altered in Atg2-PM4 mutant

Using the above described cryo-CLEM workflow with Atg2-PM4 *snf7Δ cells*, we observed a striking Atg2-PM4 phenotype with respect to phagophore morphology: 16 out of 25 phagophore rims revealed an enlarged intermembrane spacing of 50 up to 200 nm when analyzed in *z*-section (**Fig. 3a-f**). The remaining 9 rims resembled the Atg2-WT phenotype with a small widening of close-to evenly distanced juxtaposed bilayers spaced apart approximately 10 to 20 nm (**Fig. 3g-i**) as characterized previously (Bieber et al., 2022). Interestingly, when seen in cross-section, one specific phagophore had one enlarged rim region and the corresponding rim region on the other side of the same phagophore appears to be Atg2-WT-like (see **Fig. 3b**). Others exhibited either uniformly Atg2-WT-like or two enlarged rims. When inspecting membrane contact sites, as in the Atg2-WT, we also detected connecting protein densities spanning from the phagophore to putative donor membranes (**Fig. 3j** and **3h**\*). Notably, two of these tethering densities were 29 nm in length (**Table 2**), which is 9 nm longer than mammalian ATG2 and Vps13 but corresponds to the predicted length of the yeast protein Csf1, another bridge-like lipid transfer protein (Toulmay et al., 2022; Valverde et al., 2019; De et al., 2017). Another extended density of 23 nm length tethering the phagophore to the lipid droplet surface is closer to the observed length of ATG2 and Vps13 (Valverde et al., 2019; De et al., 2017). Furthermore, we also observed a direct membrane contact site of the phagophore and lipid droplet not obviously bridged by protein density. The outermost bilayer of the phagophore was found touching the phospholipid monolayer of the lipid droplet surface giving up the otherwise close-to-ideal spherical shape and leaving an increased intermembrane distance of the phagophore at the region of contact (**Fig. 3i\***). Together, this Atg2-PM4 ultrastructural analysis revealed a remarkable Atg2-PM4 phenotype of enlarged phagophore rims while they also contain typical and some thus far unobserved protein-mediated membrane contact sites between phagophore and putative donor membranes.

**Figure 3:**
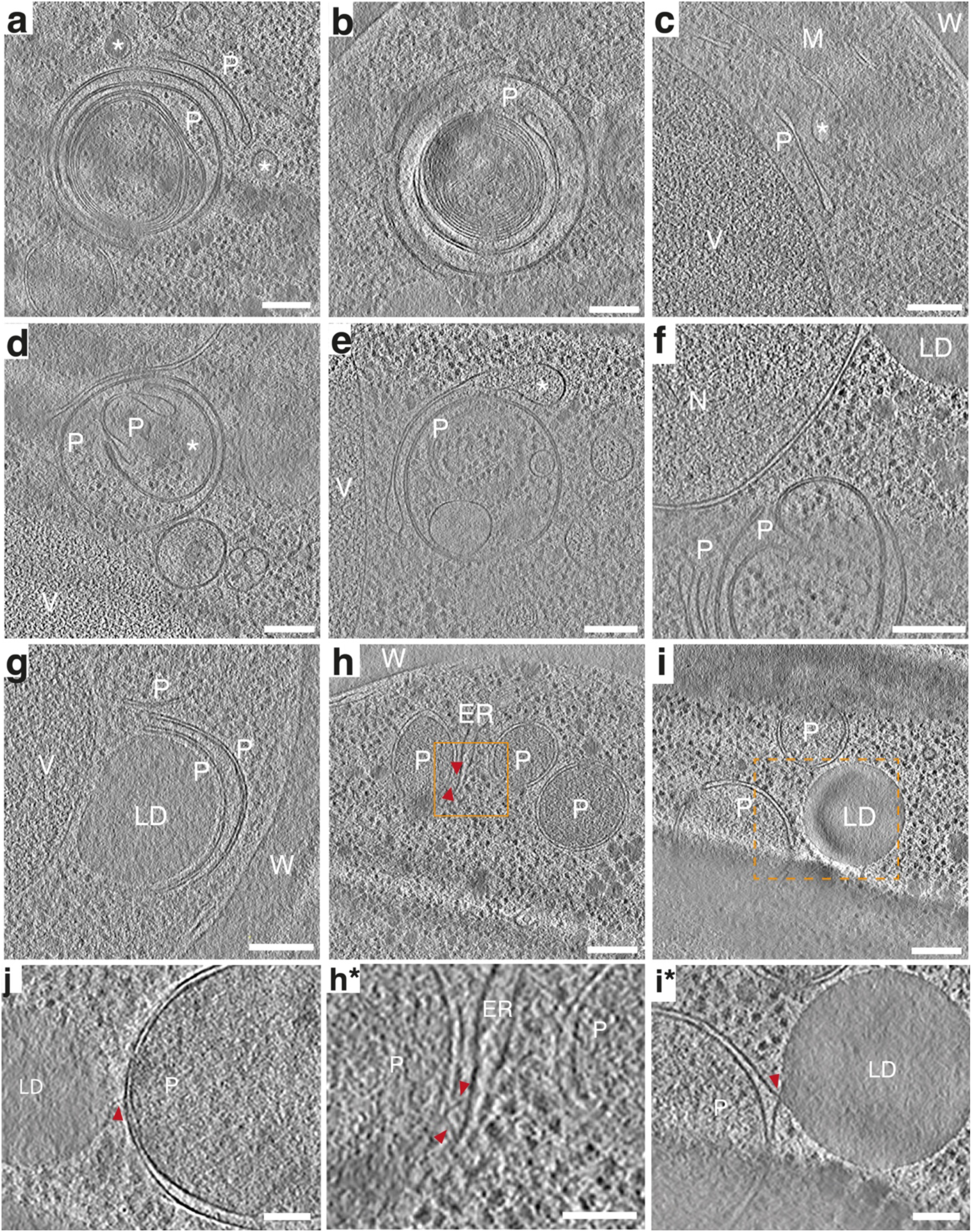
Tomogram slices of phagophore ultrastructures in Atg2-PM4 cells. **a**)-**i\***) Tomographic slices of multiple tomograms from Atg2-PM4 mutant yeast cells, denoised with IsoNet (Liu et al., 2022). Red arrows indicate elongated densities spanning from the phagophore membrane to the ER or lipid droplets, measured lengths of such densities are listed in Table 2. **h\***) Enlarged view of tomogram slice shown in panel h (orange box). **i\***) Enlarged view from a different tomographic z-slice from the tomogram shown in panel i (dashed orange box). Phagophores are indicated with P, lipid droplets with LD, endoplasmic reticulum with ER, vacuole with V, nucleus with N, and the cell wall with W. Asterisks indicate vesicles of unknown origin. Scale bars are 200 nm.

### Analysis of enlarged rims and associated membrane structures

To assess the morphological changes of the Atg2-PM4 phagophores in more detail, we specifically analyzed the phagophores from the cryo-tomograms using segmentation and grouped the Atg2-PM4 phagophore rim sections in ‘enlarged’ and ‘thin’ classes. As we analyzed representative 2D sections through the 3D phagophore we refer to the rim as tips. Subsequently, we computationally separated the tip region next to the backbone region of the phagophore making up the central portion of the phagophore (**Fig. 4a**). Comparison of membrane segmentations of the phagophore rims from representative tomographic slices of the Atg2-PM4 enlarged class showed marked differences to the Atg2-WT and the Atg2-PM4 thin classes (**Fig. 4b**). While the intermembrane distances of the backbone region of Atg2-WT, Atg2-PM4 enlarged and thin phagophore classes shared similar widths of 135 ± 25, 119 ± 29 and 135 ± 29 Å, respectively (**Fig. 4c**), the intermembrane distance at the tip (tip width) of the Atg2-PM4 enlarged class is 284 ± 91 Å as opposed to 127 ± 10 and 131 ± 26 Å of Atg2-WT and Atg2-PM4 thin, respectively (**Fig. 4d**). Similarly, the average of the maximum absolute curvatures of the Atg2-PM4 enlarged tips is significantly higher than the ones of the Atg2-WT and Atg2-PM4 thin class at 1.6 x 10^-3^ vs. 0.9 x 10^-3^ and 0.9 x 10^-3^, respectively (**Fig. 4e**). Furthermore, the Atg2-PM4 enlarged tips have a lower percentage of pixels with lower absolute curvatures and more pixels with higher absolute curvatures than the Atg2-WT and Atg2-PM4 thin phagophore tips (**Fig. 4f**). In order to understand whether the changes observed at the phagophore rim are related to the progression of phagophore growth, the rim opening angle of every tip section was estimated and related to the average tip width. The rim opening angle has been proposed as a measure of phagophore growth (Bieber et al., 2022) whereby this angle ranges from close to 0° when the phagophore is barely cup-shaped, to 180° when the tips are nearly facing each other head-on before fusing with each other (**Fig. 4g**). When plotting the rim opening angle against the mean tip width, we detected a weak linear positive trend for Atg2-WT and the Atg2-PM4-thin tips between 60 and 160° in Atg9-3x BFP cells (**Fig. 4h**), which appears to differ from previous observations proposing a negative trend in WT cells with GFP-Atg8 background (Bieber et al., 2022). The analysis of the Atg2-PM4-enlarged tip class, however, clearly reveals wider rim opening angles between 0 and 140° and widely scattered tip width measurements between 150 – 450 Å supporting no clear trend. Taken together, the detailed distance, angle, and shape analysis based on membrane segmentation confirmed that the observed morphological changes are significantly different in the Atg2-PM4 enlarged class and restricted to the phagophore rim while the phagophore backbone appears similar to the Atg2-WT.

**Figure 4:**
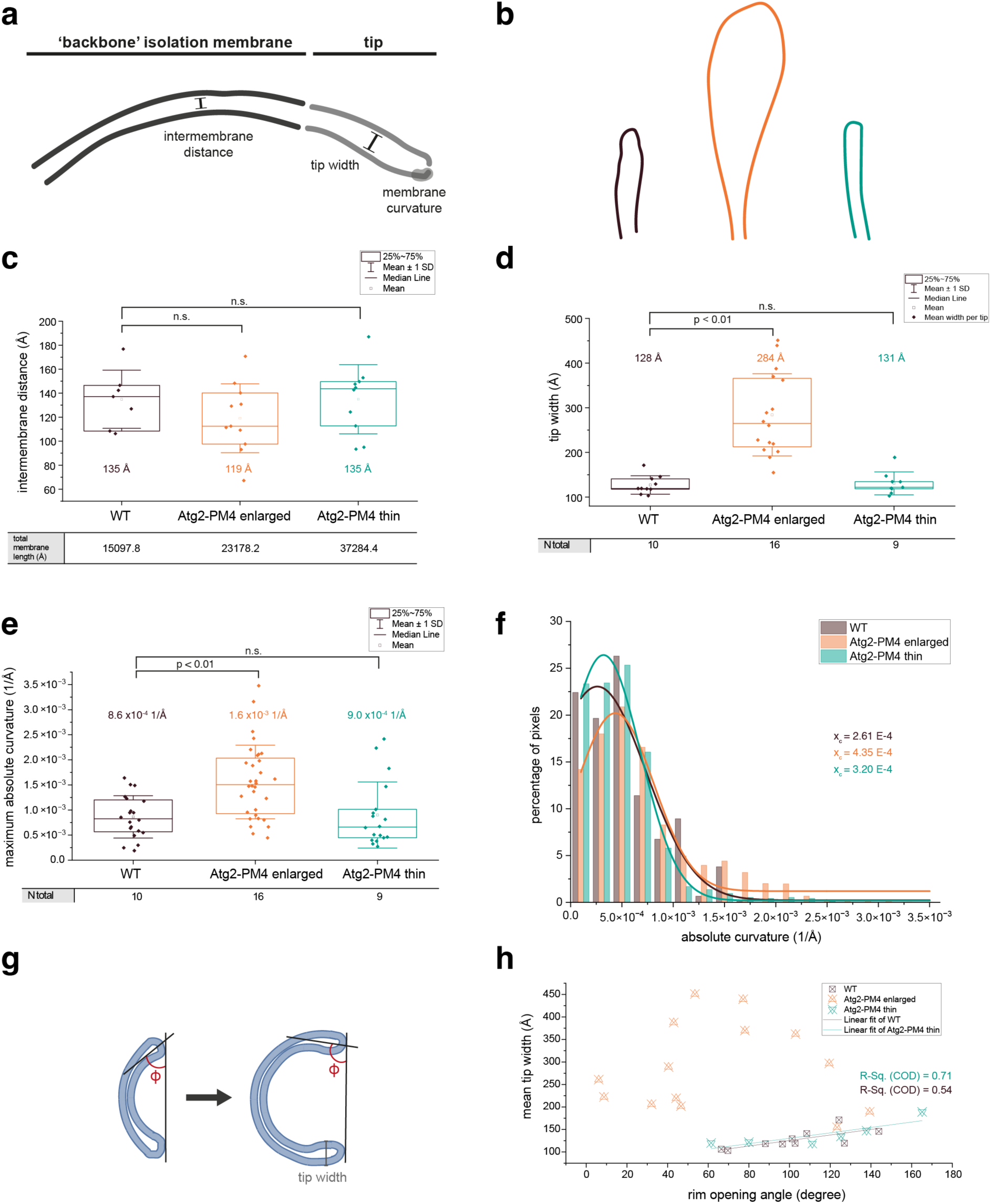
Quantification of morphological features in phagophores. **a**) Illustration of segmentation showing the analyzed parts of phagophore morphology. **b**) Representative two-dimensional segmentations of the Atg2-WT (brown), Atg2-PM4 ‘enlarged’ (orange) and Atg2-PM4 ‘thin’ (green) through phagophore rim. Tip represents the **c**) Box plot comparing the intermembrane distance of ‘backbone’ phagophore between the Atg2-WT, Atg2-PM4 enlarged and Atg2-PM4 thin. **d**) Box plot comparing mean phagophore tip width of all the segmented phagophores. **e**) Box plot comparing the maximum absolute curvature of all the segmented phagophore tips across the Atg2-WT, Atg2-PM4 enlarged and Atg2 PM4-thin phenotype. **f**) Distribution of percentage of pixels of segmented phagophore tips across different absolute curvature values with Gauss curve fits. **g**) Schematic representation of the rim opening angle as proposed by (Bieber et al., 2022). **h**) Relationship of rim opening angle and mean tip width. The R-square (coefficient of determination) and the adjusted R-square of the fitted curves are shown in the table below the graph.

When further inspecting our Atg2-PM4 tomographic data, we noticed the occurrence of vesicles in the proximity of up to 200 nm to phagophore rims (**Fig. 5a-e, Supplementary Fig. 2**). These phagophores were found in different stages of expansion varying in length and rim opening angles. Atg2-WT tomograms did not contain such vesicles. Atg2-PM4 vesicles varied in circularity and in size, with diameters ranging from 37 to 187 nm (**Table 3**). Subsequently, in one particular tomogram we also observed one phagophore with rim sections as wide as typical vesicle diameters between 150 and 190 nm suggesting a possible fusion between phagophore rims and vesicles (**Fig. 5f, 5g**). After membrane segmentation of the tomographic reconstruction displayed as a 3D isosurface rendering (**Fig. 5h**), we observed an approximately 180 nm thick slab through the membrane phagophore while we discerned two ∼200 nm wide distinct membrane entities that we believe to be two fusing vesicles attached to enlarged phagophore rims on opposite ends of the phagophore connected through a ∼100 nm wide membrane stalk. In the 3D rendering of the segmentation, the vesicles do not appear completely closed, which can be attributed to the missing wedge, a known sampling problem in electron tomography (Förster and Briegel, 2024). Taken together, in one of the tomograms, we believe to have captured cellular vesicles attached to an expanding phagophore.

**Figure 5:**
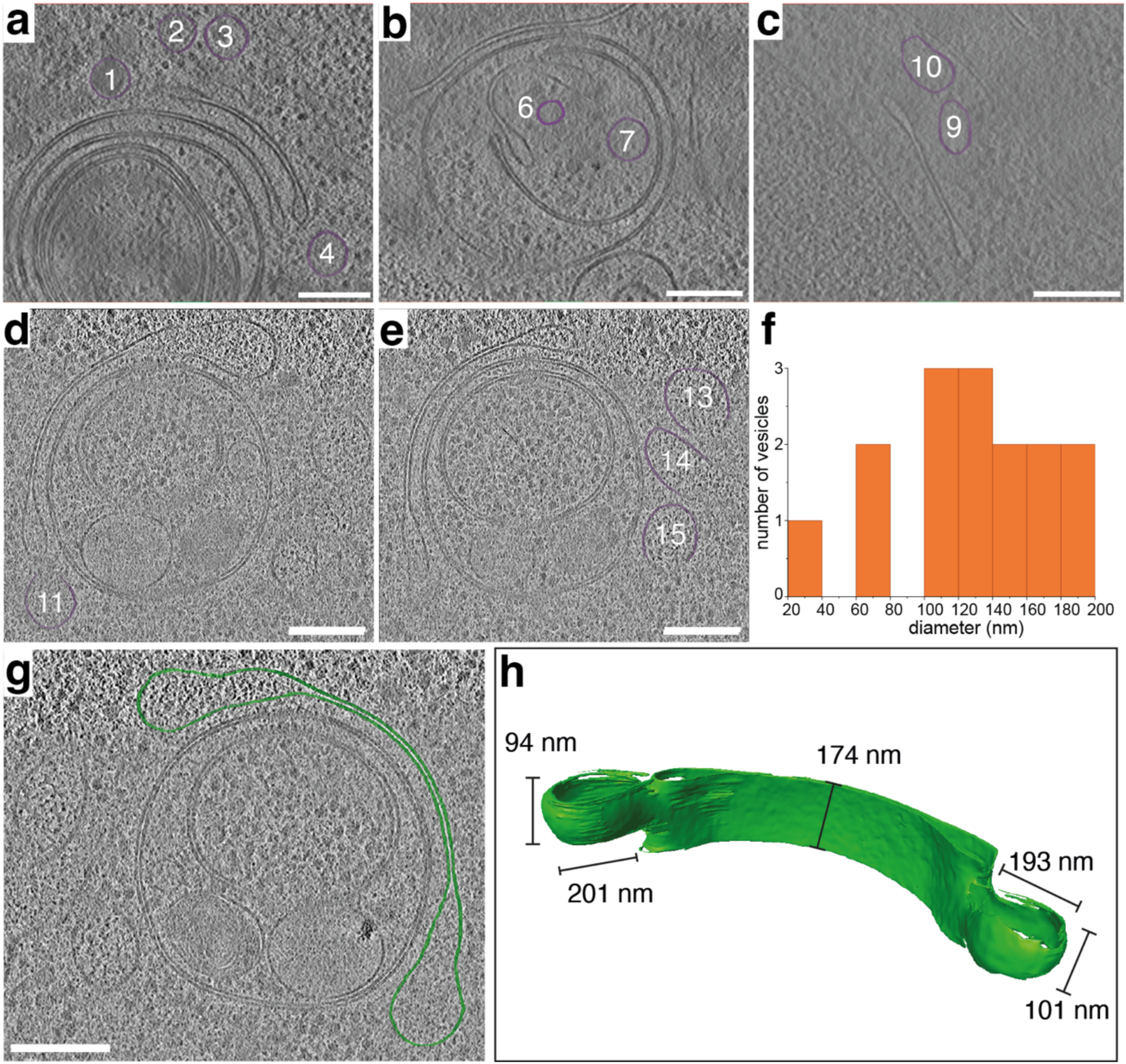
Tomographic slices of phagophore rims in proximity to cytosolic vesicles in Atg2-PM4 cells. **a**)-**e**) Vesicles proximal to phagophore rims are presented in tomographic slices. **f)** Size distribution of vesicles. Diameters and circularity values can be found in Table 3. Scale bars are 200 nm. Tomographic slices were denoised with IsoNet (Liu et al., 2022). **h**) Three-dimensional isosurface rendering of the segmented phagophore in panel g) with vesicles putatively fusing to the phagophore rims.

**Table 3:**
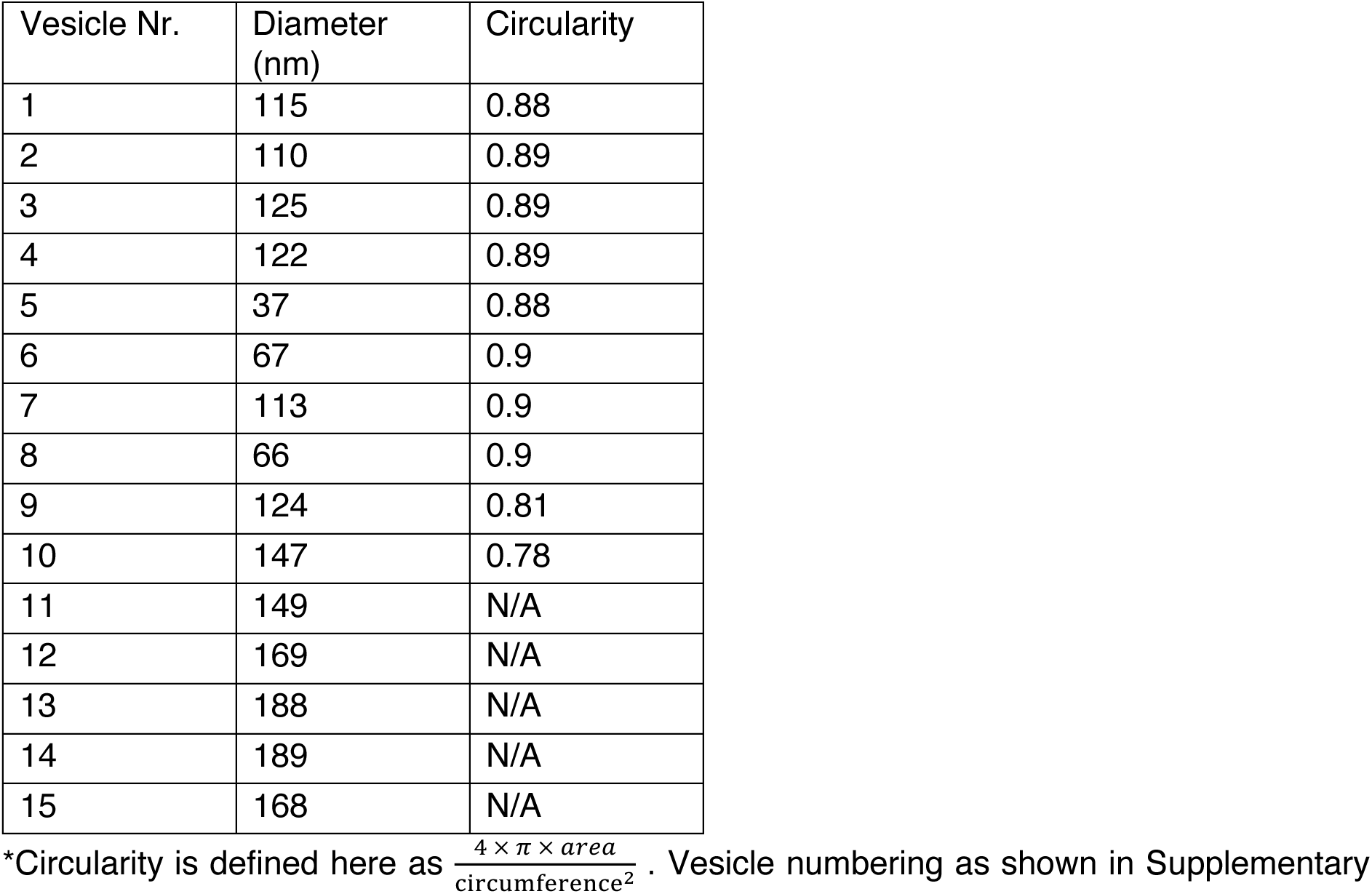
Vesicle sizes and vesicle circularity*.

As COPII coat proteins such as Sec13 and 31 have been observed by fluorescence microscopy to localize at the phagophore rims (Graef et al., 2013; Suzuki et al., 2013) and COPII vesicles have been identified as an important lipid source for autophagosome biogenesis in yeast (Shima et al., 2019), we therefore checked for the presence of characteristic protein lattice coats on the vesicles proximal to the phagophore rims (**Supplementary Fig. 2**). In these cases, we could not identify typical features of clathrin, COPII or COPI lattice coatings, such as distinct triskelion of clathrin or the triangular Sec13/31 COPII (Bykov et al., 2017; Cheng et al., 2007; Zanetti et al., 2013). Moreover, typical COP vesicles surrounding the Golgi network have been found to be around 50 nm (Bykov et al., 2017; Donohoe et al., 2013) whereas many of the here observed vesicles are larger than 100 nm. Together, these vesicular features do not allow an assignment and leave the cellular origin of the here imaged vesicles open. The observed vesicles close to the phagophore rims are unlikely a result of the *snf7Δ* strain of aberrant endosomal processing as they were not detected in Atg2 WT cells (see **Fig. 1d, e**), but rather belong to membranous structures of different subcellular identity. Taken together, the presence of medium-sized 40-200 nm vesicles close to the phagophore rims and in some cases enlarged rims of the same size suggest vesicle fusion to the rims of the phagophore in Atg2-PM4 cells.

## Discussion

Using *in situ* cryo-ET guided by cryo-CLEM, we imaged and analyzed the natively preserved morphology of expanding phagophores of *snf7Δ S. cerevisiae* strains. We observed multiple phagophore-membrane contact sites including direct lipid droplet contact and linkages mediated by bridge-like proteins. Using a previously characterized Atg2-PM4-mutant unable to interact with Atg9 (Gómez-Sánchez et al., 2018), we found an altered phagophore morphology. The Atg2-PM4 mutant revealed an about two-fold enlarged phagophore rim width while it also showed a significantly decelerated autophagosome biogenesis in live-cell imaging. As we observed the presence of 40 to 200 nm large vesicles in close proximity to phagophore rims and captured vesicles attached to a phagophore, we reason that the enlarged rims in our tomographic data represent an intermediate structure resulting from vesicular fusion to the phagophore rims in a lipid-transport impaired mutant.

The disruption of the Atg2-Atg9 interaction caused by the alanine substitution in positions 1264-1268 in the Atg2-PM4 mutant clearly affected the observed phagophore morphology leading to enlarged phagophore rims. Due to the disrupted interaction with Atg9, Atg2 was found mislocalized across the entire phagophore in contrast to Atg9 and Atg18 that remained localized at the phagophore rims (Gómez-Sánchez et al., 2018). As a consequence of the disrupted Atg9 interaction by another mutant, a reduction of the human Atg2 *in vitro* lipid transfer activity has been reported (Van Vliet et al., 2022). Conversely, *in vitro* Atg2 lipid transfer was shown to increase in the presence of Atg18 and PI3P (Osawa et al., 2019). Therefore, the disruption of described Atg2 interactors is likely to have an overall attenuating effect on Atg2-mediated lipid supply to the phagophore. Given the close spatial coupling and proposed lipid channeling of Atg2 with Faa1 and Atg9 (Schütter et al., 2020; Van Vliet et al., 2022; Wang et al., 2025), the effective lipid transport and subsequent lipid equilibration across the leaflets is likely to be altered in the Atg2-PM4 mutant. Under wildtype conditions, due to close spatial coupling Atg2 provides lipid substrates unidirectionally for the Atg9 scramblase (**Fig. 6a**). In the Atg2-PM4 mutant, the lack of unidirectional lipid substrate provision for Atg9 may affect the lipid turnover (**Fig. 6b** left) possibly leading to changes in the rim shape considering principal membrane geometry arguments (Frolov et al., 2011). A recent study tested a closely related Atg9-binding mutant, known as Atg2-PM1 that showed equal mislocalization of Atg2 and failed to perform rim aperture constriction also possibly due to changes in membrane geometry (Gómez-Sánchez et al., 2018; Shatz et al., 2024). Similarly, in our Atg2-PM4 mutant, we observed a disrupted correlation of mean rim width and rim opening angle of the Atg2-PM4 enlarged rims with respect to the wildtype-like rims. In line with a series of prior studies, our investigation further supports the requirement of precise temporal and spatial localization of Atg9-Atg2-Atg18 complexes at the phagophore for autophagosome biogenesis. The observation of a putative vesicle-phagophore fusion event to an expanding phagophore in cells with decelerated autophagosome biogenesis when compared with WT cells may arise from two possible scenarios. First, in Atg2-WT cells the fusion event may be too transient to be captured by our cryo-CLEM approach while it is slower and becomes detectable in the Atg2-PM4 mutant. Second, the Atg2-PM4 mutation is likely to attenuate the primary lipid supply mechanism to such an extent that the cell compensates by engaging alternative lipid delivery pathways to the phagophore (**Fig. 6b** right).

**Figure 6:**
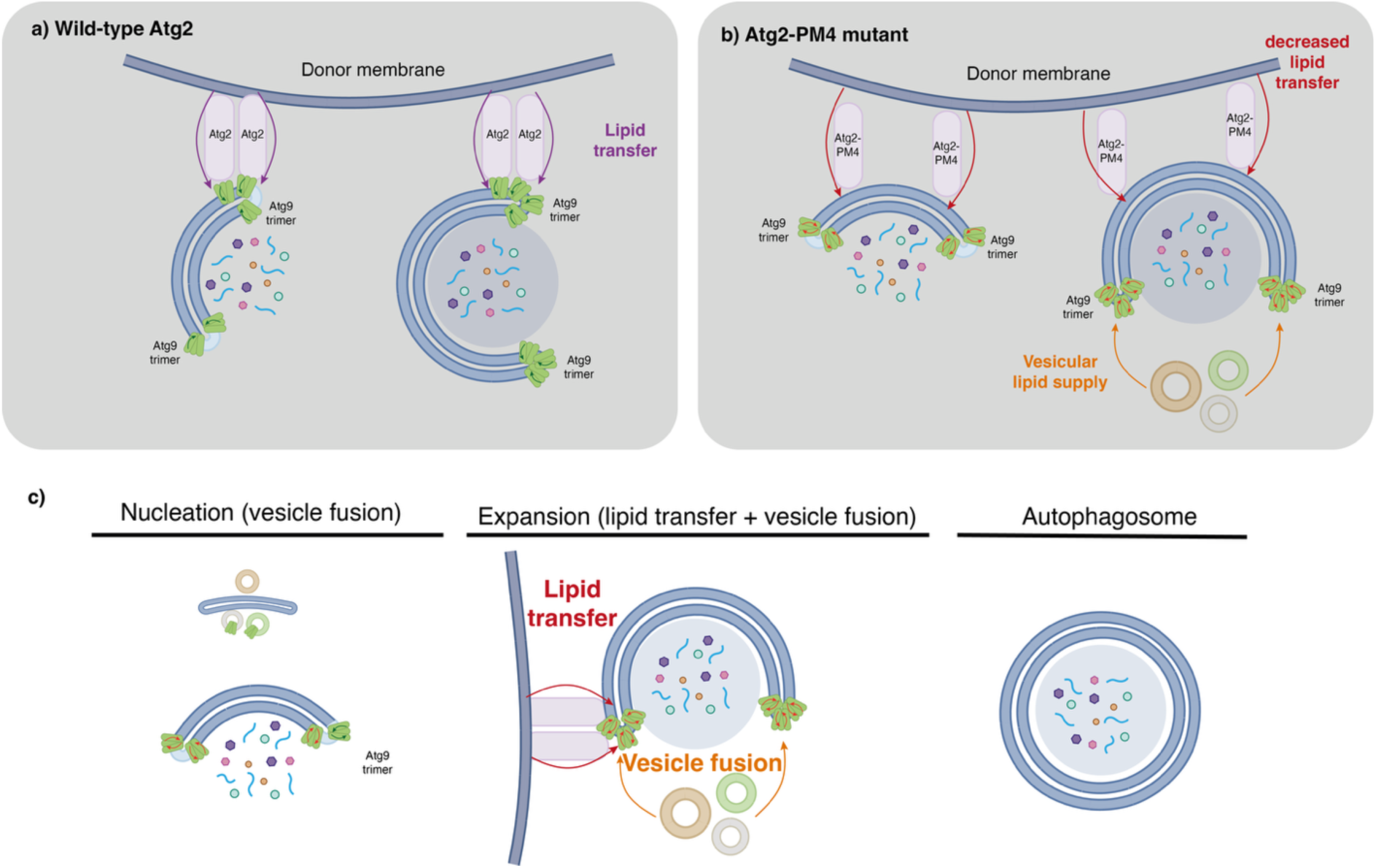
Vesicle fusion can contribute to phagophore expansion. **a**) Current phagophore expansion model in wild-type Atg2 cells. Atg2-WT spatially coupled with Atg9 is localized to the phagophore rim and transfers lipids (purple arrows) from the donor membrane to the phagophore rim. Atg9 transfers lipids to the inner leaflet of the phagophore membrane by its scramblase activity (dark green arrows). As a result of the Atg2-Atg9 lipid transfer-scramblase activities, the phagophore grows. **b**) Proposed model of delayed phagophore expansion activity in Atg2-PM4 cells: through mislocalization of Atg2 (large red arrows) and decoupled Atg9 scramblase activity (small red arrows) growth of phagophore membrane is delayed. Cellular vesicles can provide additional lipids for its expansion (orange arrows). **c**) During autophagosome biogenesis, small Atg9 vesicles fuse at the phagophore assembly site during early stages to form a nascent phagophore. Once the phagophore has established the corresponding membrane contact sites with the ER/ERES, the phagophore membrane expands by lipid transfer from the donor membrane and a possible contribution of vesicle fusion at the phagophore rims.

Interestingly, the cellular origin of the observed vesicles could not be fully clarified. Apart from the established phagophore vicinity to the ER and lipid droplets, we could not identify any other cellular compartment associated with the vesicles. In the absence of experimental support, it is tempting to speculate that the observed vesicles may be identical to the earlier characterized Golgi-derived Atg9 vesicles (Yamamoto et al., 2012). However, most of the vesicles visualized here in cells were about 90 – 180 nm in diameter, i.e. two to three-fold larger than the previously reported size of 30 – 60 nm of isolated Atg9 vesicles (Yamamoto et al., 2012). These smaller vesicles were also shown to occur in clusters and tubules at the initial stages of autophagy (Mari et al., 2010) not at an expanding phagophore as visualized here (**Fig. 6c**).

Our live-cell imaging data confirmed that the observed vesicles are not class E compartment derived and therefore not an artifact from the *snf7Δ* strain. Furthermore, our data are consistent with previous reports showing that Atg8 clusters do not overlap with class E compartments in *snf7Δ* as well as *vps4Δ* ESCRT mutants, under both growing and nitrogen-starvation conditions (Zhou et al., 2019). The recent observation of membrane contact sites between the phagophore rim and the ERES by Atg2 and TRAPPIII complex (Gómez-Sánchez et al., 2025) may explain why COPII vesicles have been found colocalized with phagophore rims and identified as a membrane source previously (Graef et al., 2013; Shima et al., 2019; Suzuki et al., 2013). The diameters of canonical COPII vesicles were characterized in the size range 60 – 80 nm (Kim et al., 2005) while they can also be significantly larger up to 300 nm in the presence of cargo (Gorur et al., 2017). The resolution and available details in the yeast tomograms were not sufficient to assign the nature of the vesicles neither for COPI nor clathrin lattices. While the images reveal the vesicles attached to the phagophore rims rather than the back of the phagophore, it remains open what the specific mechanisms behind vesicle trafficking to the phagophores and signals for vesicle-phagophore fusion are.

Regardless of our most striking observations on the phagophore morphology, we would also like to emphasize that about one third of phagophore rims in the Atg2-PM4 phenotype had a width resembling the Atg2-WT, which could be explained by the presence of alternative lipid transporters such as Vps13 or Csf1 (Dabrowski et al., 2023; Toulmay et al., 2022). Indeed, Vps13 has been shown to be a lipid transporter acting complementarily and non-rate limiting to Atg2 in autophagy (Dabrowski et al., 2023) possibly coordinated by a direct Atg9 interaction (Van Vliet et al., 2024). Csf1 has a putative length of 30 nm, which is consistent with the estimated lengths of densities observed tethering a phagophore with the ER close to the phagophore rim (see **Fig. 3h*, Table 2**). Moreover, it has recently been shown that Csf1 localizes at ER-plasma membrane and ER-mitochondrial contact sites (Toulmay et al., 2022). Furthermore, Csf1 has been suggested to act as a lipid transfer protein in phosphatidyl ethanolamine metabolism and lipid homeostasis (Toulmay et al., 2022; John Peter et al., 2022). The occurrence of wildtype-like phagophore rims suggests that other lipid-transfer mechanisms exist that can take over some of the tasks typically performed by Atg2. Interestingly, we also observed a direct contact between lipid droplets and the phagophore. To our knowledge, the direct transfer of phospholipids from the phospholipid monolayer on the surface of a lipid droplet to the phagophore in this way has not been observed so far by electron microscopy. Further studies are required to disentangle the contributions of the multiple players, Atg2, Vps13, Atg9 and Atg18 involved in the delivery of lipids to the growing phagophore.

Our tomographic observations suggest that vesicles are fusing with the rims of the phagophore rather than the sheet-like phagophore backbone. Given the high membrane curvature present at the phagophore rims, this geometry is likely to be favored for membrane fusion with spherical cellular vesicles. According to EM observations on the Golgi-apparatus, vesicles were also shown to primarily fuse with the rims rather the sheets (Donohoe et al., 2013). Together, our cryo-ET data recorded *in situ* reveal vesicle fusion at the stage of expanding phagophores and adds to the cellular repertoire of processes contributing to phagophore expansion.

## Materials and methods

### Yeast strains

The yeast strains used in this study are listed in **Supplementary Table 1**. Genomic deletions were characterized with respect to expression and autophagic flux as shown in **Supplementary Fig. 4**. Yeast genomic insertions and tagging were performed according to Janke et al. (Janke et al., 2004), and multiple deletions were generated by PCR knockout, mating, and dissection. GFP-ATG8-containing strains were generated by crossing with yTB282 (*hi3Δ1 leu2Δ0 met15Δ ura3Δ0 atg8*::GFP-*ATG8*), which had been generated by seamless tagging (Khmelinskii et al., 2011). The genotype of generated strain was confirmed by colony PCR and Western blot analysis.

For live-cell imaging of Vps8 and Atg8, the *ATG2* coding region was replaced with the *hygromycin-resistance* gene *(hphNT1)* to create the atg2*Δ* strain, using PCR primers containing 60 bases of identity to the regions flanking the open reading frame (Janke et al., 2004). Gene knockout was verified by polymerase chain reaction (PCR) and Ape1 processing analysis (Mari et al., 2010). PCR-based integration of *mScarlet* tag at the 3’ end of *VPS8* was used to generate a strain expressing C-terminal fusion protein under the control of the authentic promoter. The plasmid template for integration was pFA6-mScarlet-NatMX6.

For live-cell imaging of Atg2 and Atg8, *Δatg2::atg2^PM4^-3xGFP-CaURA3* strains were generated by assembling the following fragments via gap repair into a pRS315 plasmid backbone: (1) a 1000 bp long region 5’-upstream of the endogenous *ATG2* locus, (2) full length *atg2^PM4^* C-terminally tagged with *3xGFP-CaURA3* derived from a pFA6a*-link-3yEGFP-CaURA3 plasmid*, and (4) a 1000 bp long region 3’-downstream of the endogenous *ATG2* locus. Following isolation, the plasmid was sequenced and enzymatically digested using the restriction enzymes *Spe*I and *Sal*I. Next, the cut plasmid fragment was transformed into pRS304-mCherry*-ATG8 Δatg19::His3MX6 Δatg2::NatMX6* cells. The Δ*atg2::Nat3MX6* cassette was replaced with the *atg2^PM4^-3xGFP-CaUraMX6* construct via homologous recombination at the endogenous *ATG2* locus, resulting in the generation of pRS304-mCherry*-ATG8 Δatg19::His3MX6 Δatg2::atg2^PM4^-3xGFP-CaURA3* cells.

### Growth conditions

Strains were grown in rich (YPD) media containing 1 % yeast extract, 2 % peptone (BD) and 2 % glucose; or in synthetic drop-out media (-Leu,-Ura, or-Leu-Ura, Formedium) supplemented with 2 % glucose; or nitrogen starvation medium (SD-N) media containing 0.17 % yeast nitrogen base without amino acids or ammonium sulfate (ThermoFisher Scientific) and 2 % glucose to mid-log phase. Pre-cultures (10 ml) for vitrification in either YPD or synthetic drop-out media were inoculated with a single colony. Cells were cultured at 30°C for at least 14 hours to ensure cells would not be stressed. The O.D._600_ was monitored and as soon as it was between 0.55 and 0.8 the culture was used for starvation and plunge freezing. For nitrogen starvation, the cells were pelleted at 3000 x *g* for 5 minutes and washed three times with SD-N media. Cells were incubated in SD-N media for a minimum of 2 hours and a maximum of 5 hours before vitrification.

### Yeast cell transformation

Plasmids used in this work are listed in **Supplementary Table 2**. Transformation procedures are described in (Gietz, 2014) as the’Quick and Easy Transformation Protocol’ with minor modifications also known as the LiAc/SS Carrier DNA/PEG method. In brief, the transformation buffer was prepared in advance with 0.24 M Lithium acetate, 47% PEG 3350 (sterile filtered). The reaction mix was composed of 5 μl single stranded salmon sperm carrier DNA (Sigma Aldrich), 85 μl transformation buffer, 10 μl 1M DTT and 0.5 to 2 μl of the plasmid (300 to 500 ng/μl concentration). The yeast patch was scraped off the agar with a sterile loop and mixed into the transformation reaction mix directly. This suspension was very briefly vortexed and incubated at 45°C for exactly 30 min. Finally, the reaction mix was plated out on the appropriate selection agar plate.

### Detection of expression and autophagy flux

Yeast cell cultures were precipitated with 7% trichloroacetic acid (TCA) for 30 min on ice. Precipitated proteins were pelleted at 16,000 x *g* for 15 min at 4°C, washed with 1 ml acetone, air-dried, resuspended in urea loading buffer (120 mM Tris-HCl pH 6.8, 5% glycerol, 8 M urea, 143 mM β-mercaptoethanol, 8% SDS), boiled and analyzed by SDS-PAGE followed by immunoblotting.

### Antibodies

The following antibodies were used in this study: rabbit polyclonal anti-Snf7 (1:5,000, David Teis, Medical University Innsbruck), rabbit polyclonal anti-Ape1 (1:15,000), was generated by immunizing rabbits with a synthetic peptide corresponding to amino acids 168-182 (Papinski et al., 2014), mouse monoclonal anti-RFP (1:1,000, 6g6, Ref No. 6g6-100 Lot No51020014AB-05, Chromotek).

### Live-cell fluorescence microscopy

Imaging of Vps8 and Atg8: To examine bulk autophagy, cells were grown in SD-URA medium before transferring them in SD-N medium for 2 hours. Cells were incubated with CellTracker™ Blue 7-amino-4-chloromethylcoumarin dye (CMAC, ThermoFisher Scientific, C2110) as previously described (Stefan and Blumer, 1999) for 10 minutes to stain the vacuolar lumen. Images were acquired with a DeltaVision Elite RT microscope system (GE Healthcare, Applied Precision), equipped with a UPLASPO 100× oil/1.40 NA objective, a pco.edge 5.5 sCMOS camera (PCO) and a seven-colour InsightSSI solid-state illumination system (GE Healthcare, Applied Precision). Images were generated by collecting a Z-stack of 22 pictures with focal planes 0.20 μm apart to cover the entire volume of a yeast cell, and subsequently deconvolved using the SoftWoRx software (Applied Precision). A single focal plane is shown in the figures. Percentage of Atg8-positive structures associated to the Vps8 clusters was determined by analyzing 50 cells from three independent experiments. Data represent the average of three independent biological replicates ± standard deviation (SD). The shown data are those of a representative experiment. Data were statistically analyzed with GraphPad Prism 6 (GraphPad Software Inc.), using a two-tailed unpaired *t*-test with Welch’s correction. All comparisons with a *p*-value less than 0.05 (*p* < 0.05) were considered statistically significant and indicated with a single asterisk.

Imaging of Atg2 and Atg8: *S. cerevisiae* cells were imaged at room temperature after 1 hour of nitrogen starvation in 96-well glass-bottom microplates (Greiner Bio-One). Live-cell images and semi-three-dimensional time-lapse fluorescence images were taken using a Dragonfly 500 spinning disk microscope (Andor) attached to an inverted Ti2 microscope stand (Nikon) with a CFI Plan Apo Lambda 60Å∼/1.4 oil immersion objective (Nikon) and a Zyla 4.2 sCMOS camera (Andor). Following acquisition, images were processed using Fiji version 2.1.0. The number of Atg8 puncta and expanded autophagosomal membranes including expanded tubular and ring-shaped structures was analyzed after 1 hour of nitrogen starvation, by randomly marking 50 cells with the Fiji multipoint tool.

### Cell vitrification

Nitrogen starved *S. cerevisiae* cells were applied to Quantifoil (Großlöbichau, Germany) grids (Cu 200 mesh, R2/1, either Carbon or SiO2 film). In some cases, carboxylate-modified microspheres (1 μm diameter crimson or 1 μm diameter blue, Invitrogen, ThermoFisher Scientific, Waltham, MA, US) were added to the cell suspension prior to plunge freezing. Cells were vitrified using a liquid ethane/propane mixture or liquid ethane at liquid nitrogen temperature and a Vitrobot Mark IV (ThermoFisher Scientific, Waltham, MA, USA). Back-side blotting was carried out by covering the paddle of the device on the side of the grid film with a Teflon disc (generous donation from Dr. Julia Mahamid). The paddle at the back side of the grid was covered with filter paper (Whatman 597, Maidstone, UK) as customary. For a given grid, 7 μl of cell suspension were applied and after one second wait time excess liquid was blotted away. The blot time ranged from 10 to 13 seconds. Blot force was kept consistently at-10, chamber humidity at 80 % and chamber temperature at 30°C. After plunge-freezing the grids, they were usually clipped into autogrids (ThermoFisher Scientifc, Waltham, MA, USA) for further electron microscopy applications.

### Lamella Preparation

For this work, the Aquilos 2 dual-beam FIB microscope (ThermoFisher Scientific, Waltham, MA, USA) was used. Grids were kept at approx.-195 °C during the entire milling procedure. FIB-milling procedures were carried out as for instance described in (Schaffer et al., 2017) with some modifications. Lamella were milled using the gallium beam at 30 kV (0.1 nA to 10 pA) with a typical milling angle of 10 ° (i.e. stage angle of 17 °). After setting eucentric height at the sites of interest, the grids were sputter-coated with platinum (30 mA, 0.1 mbar) for 15 seconds, then coated with an organometallic platinum layer using the gas injection system (GIS) for 90 to 105 seconds. Subsequently, another sputter-coating layer was applied (30 mA, 0.1 mbar) for 15 seconds. Most of the milling sessions were carried out using the automated milling software AutoTEM (ThermoFisher Scientific, Waltham, MA, USA), followed by manual polishing. Once the lamellae were at their final thickness, in some cases a final sputter coat would be applied (30 mA, 0.1 mbar) for three seconds. Lamellae were imaged on the retro-fitted Meteor (Delmic BV, Delft, NL) bright-field cryo-fluorescence microscope with 50x 0.5 NA (Olympus LMPLFLN WD 10.6 mm) objective. The sample was kept at approx.-195 °C on the stage of the Aquilos 2 during measurements. Except for one tomogram (**Fig. 1i**), all tomographic data was collected based on FM-TEM correlation from images collected with the Meteor.

### Cryo-electron tomography

After preparation and fluorescent imaging of the lamellae, the grids were transferred to the following transmission electron microscopes: a Talos Arctica (200keV, ThermoFisher Scientific, Waltham, MA, USA) microscope together with a K3 direct electron detector with Quantum GIF filter (Gatan Inc., Pleasanton, CA, USA) and Titan Krios (300 keV, ThermoFisher Scientific, Waltham, MA, USA) equipped with a Falcon 4 direct electron detector (ThermoFisher Scientific, Waltham, MA, USA). In both cases, cameras were operated in movie mode with 10 to 15 frames per movie. For data collection and correlation of TEM overview images with fluorescence data on the Talos Arctica, the software package SerialEM (Mastronarde, 2003) was used. For acquisition of lamella overview images and correlation with fluorescence data on the Titan Krios the Maps software (ThermoFisher Scientific, Waltham, MA, USA) was utilized. For automated acquisition of tilt series on the Titan Krios the Tomography software (ThermoFisher Scientific, Waltham, MA, USA) was used. Tilt series were acquired using the dose-symmetric acquisition scheme (Hagen et al., 2017) implemented in both SerialEM and Tomography over an angular range of-60° to +60° with 2° increments. The set defocus was-5 μm and cumulative dose was kept below 130 e^-^/Å. The pixel size ranged from 1.74 to 2.78 Å depending on the camera and camera operation mode. Only four tilt series were collected on the Titan Krios, these were tilt series of the Atg2-WT of which two tomographic slices are shown in Fig. 1g and h. All other tomographic data analyzed in this manuscript were acquired on the Talos Arctica.

### Tomographic image processing

For the tomographic data processing and analysis workflow, acquired tilt frames were preprocessed using WARP (Tegunov and Cramer, 2019). Patch-tracking guided tilt series alignment and tomogram reconstruction based on weighted back projection was done with IMOD (Mastronarde, 1997; Kremer et al., 1996) or AreTomo (Zheng et al., 2022). Denoising and missing wedge correction were carried out with the help of IsoNet (Liu et al., 2022). Segmentation was done with MemBrain v2 (Lamm et al., 2024) (an updated version from MemBrain (Lamm et al., 2022)) using its pretrained model (Version b from 10th Aug 2023) on the non-denoised data. Segmentations were subsequently manually corrected and completed using Dragonfly software, Version 2022.2 (Comet Technologies Canada Inc.).

The data evaluation and quantification of the presented tomograms were carried out on tomographic slices in 2D with the help of CryoVIA (Schönnenbeck et al., 2025). Segmentations of regions of interest were manually separated, saved to individual.tif files and input into the toolkit. Prior to curvature measurements, a Gaussian filter (window size of 9 and sigma of 4) was applied in order to avoid overfitting curvature inaccuracies due to small errors in the segmentation. Subsequently a single pixel-wide skeleton of the membrane was generated. For each pixel of this skeleton, the curvature was determined by selecting the neighboring pixels in each direction along the skeleton up to 500 Å away from the pixel of interest. Next, a circle with the best possible fit through these pixels was generated. The inverse of this circle’s radius was defined as the curvature of the pixel of interest. For distance measurements, the single pixel skeleton was also used. A support vector machine was used to determine a line separating the two sides of the segmented rim sections, i.e. tips. The length of perpendicular lines (relative to the separation line) connecting individual pixels of each side of the tip were then determined and defined as the distance between the given two pixels. Circularity is defined here as 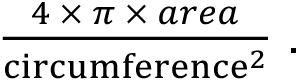 Circularity of vesicles could only be determined for vesicles where the segmentation was complete and had no gaps, as the area was otherwise not clearly defined. Data analysis and plotting were carried out in Origin(Pro) 2023 (Version 10.0.0.154, OriginLab Corporation, Northampton, MA, USA).

### Illustrations and figures

Fluorescence images from the Meteor bright-field cryo-fluorescence microscope were generated using the proprietary software Odemis (Delmic BV, Delft, NL). Figures of tomograms were made using IMOD (Kremer et al., 1996) and Dragonfly software, Version 2022.2 (Comet Technologies Canada Inc.). For live-cell imaging figure preparation, Fiji, Photoshop CC and Illustrator CC software (Adobe) were used. Some of the structural biology software used in this work, such as IMOD, AreTomo, IsoNet, and PyMOL was compiled and configured by SBGrid (Morin et al., 2013).

## Data availability

Tomography data is available at the EMDB, accession number XXX. The remaining data is available upon request to the corresponding author.

## Acknowledgements

We gratefully acknowledge the electron microscopy training, imaging and access time granted by the life science EM facility of the Ernst-Ruska Centre at Forschungszentrum Jülich. We thank Dr. Julio Ortiz for SerialEM training and support in setting up the SerialEM imaging settings. We also thank David Kartte for his help in setting up processing software and segmentation training. We thank Prof. Fulvio Reggiori (Department of Biomedicine, Aarhus University, Denmark) for reagents. The authors gratefully acknowledge the computing time granted by the JARA Vergabegremium and provided on the JARA Partition part of the supercomputer JURECA at Forschungszentrum Jülich (Thörnig, 2021). The authors acknowledge financial support through Gravitation grant “FLOW” (024.006.036) from the Dutch Ministry of Education, Culture, and Science (OCW) (R. G.-S.). The Kraft laboratory has received funding from the Deutsche Forschungsgemeinschaft (DFG, German Research Foundation) Project-ID 450216812, Project ID 409673687, SFB 1381 (Project ID 403222702), under Germany’s Excellence Strategy (CIBSS - EXC-2189-Project ID 390939984), from the European Research Council (ERC) under the European Union’s Horizon 2020 research and innovation programme under grant agreement No 769065. This work reflects only the authors’ view and the European Union’s Horizon 2020 research and innovation programme is not responsible for any use that may be made of the information it contains.

## Author contributions

C. O. P. N.: Conceptualization, EM sample preparation, EM data acquisition, EM data analysis, and manuscript preparation (original draft, review, and editing). M. L.: Generation of yeast strains, biochemical and fluorescence microscopy validation, conceptualization and scientific discussions, and manuscript review. R. D.: Generation of yeast strains, fluorescence microscopy, investigation, scientific discussions, and manuscript review. R. G.-S.: Generation of plasmids and yeast strain, fluorescence microscopy, investigation, scientific discussions, and manuscript review. S. B.: Training and technical support in CLEM sample preparation and EM data acquisition, scientific discussions, manuscript review. P. S.: Coding and running of evaluation software, and manuscript review. M. G.: Investigation, resources, scientific discussions, and manuscript review. C. K.: Investigation, resources, scientific discussions, and manuscript review and editing. C. S.: Conceptualization, investigation, funding acquisition, project administration, resources, supervision, and manuscript preparation (original draft, review, and editing).

---

**Supplementary Figure 1:**
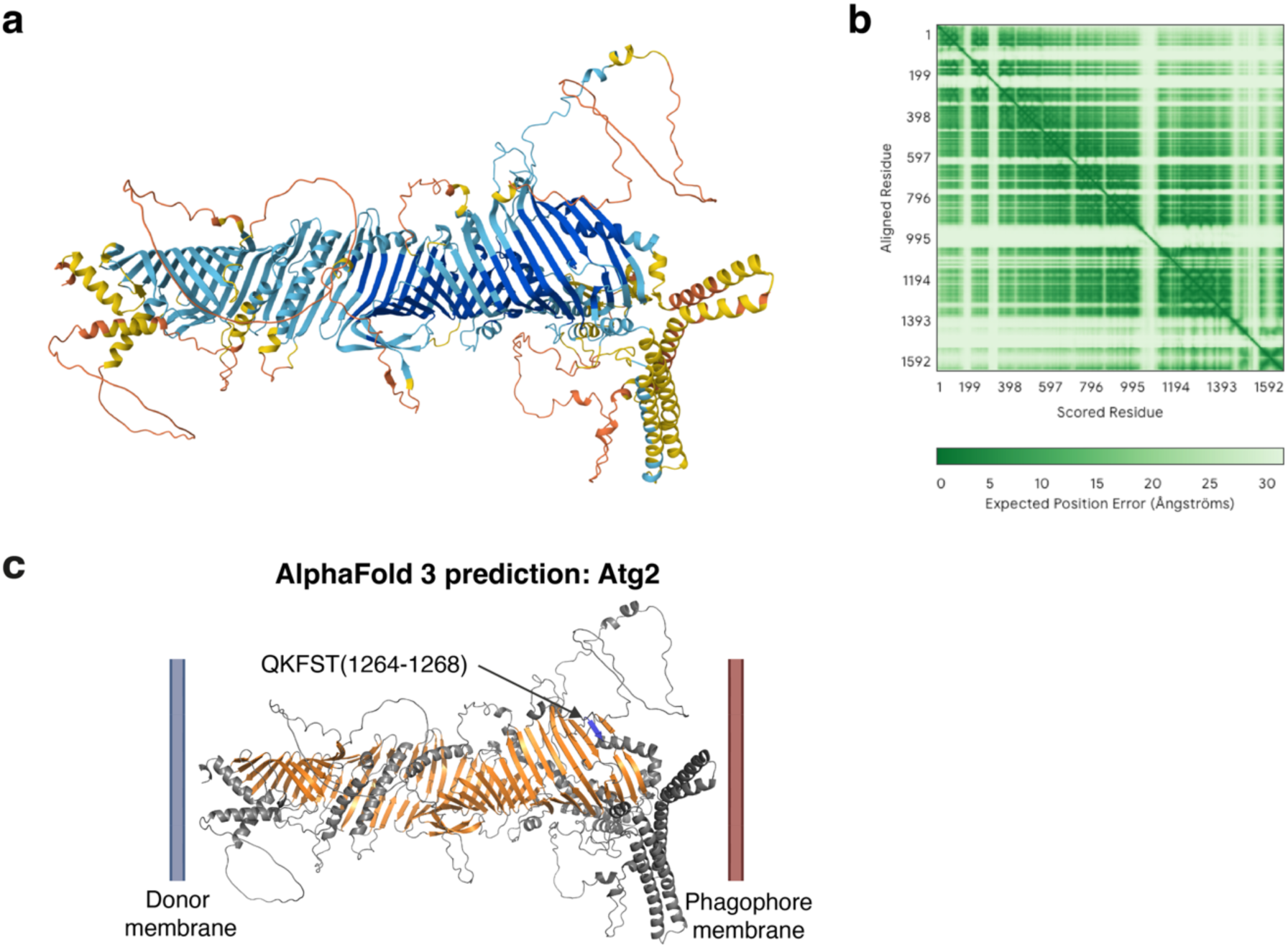
AlphaFold 3 prediction of *S. cerevisiae* Atg2. **a**) Predicted model shown in ribbon representation with pLDDT (predicted local distance difference test) score for each residue shown with the following color code; pLDDT > 90 dark blue, pLDDT > 70 light blue, pLDDT > 50 yellow, pLDDT < 50 orange. Prediction and illustration were generated using the AlphaFold 3 server with the protein sequence from UniProt (ID: P53855). **b**) PAE (predicted aligned error) score plot as generated by the AlphaFold 3 server (Abramson et al., 2024). **c**) The residues mutated in the Atg2-PM4 AF3 model are highlighted in blue. The β-sheets characteristic for the bridge-like lipid transfer protein family are shown in orange. Donor and phagophore membrane are illustrated at the N and C-terminus of Atg2, respectively.

**Supplementary Figure 2:**
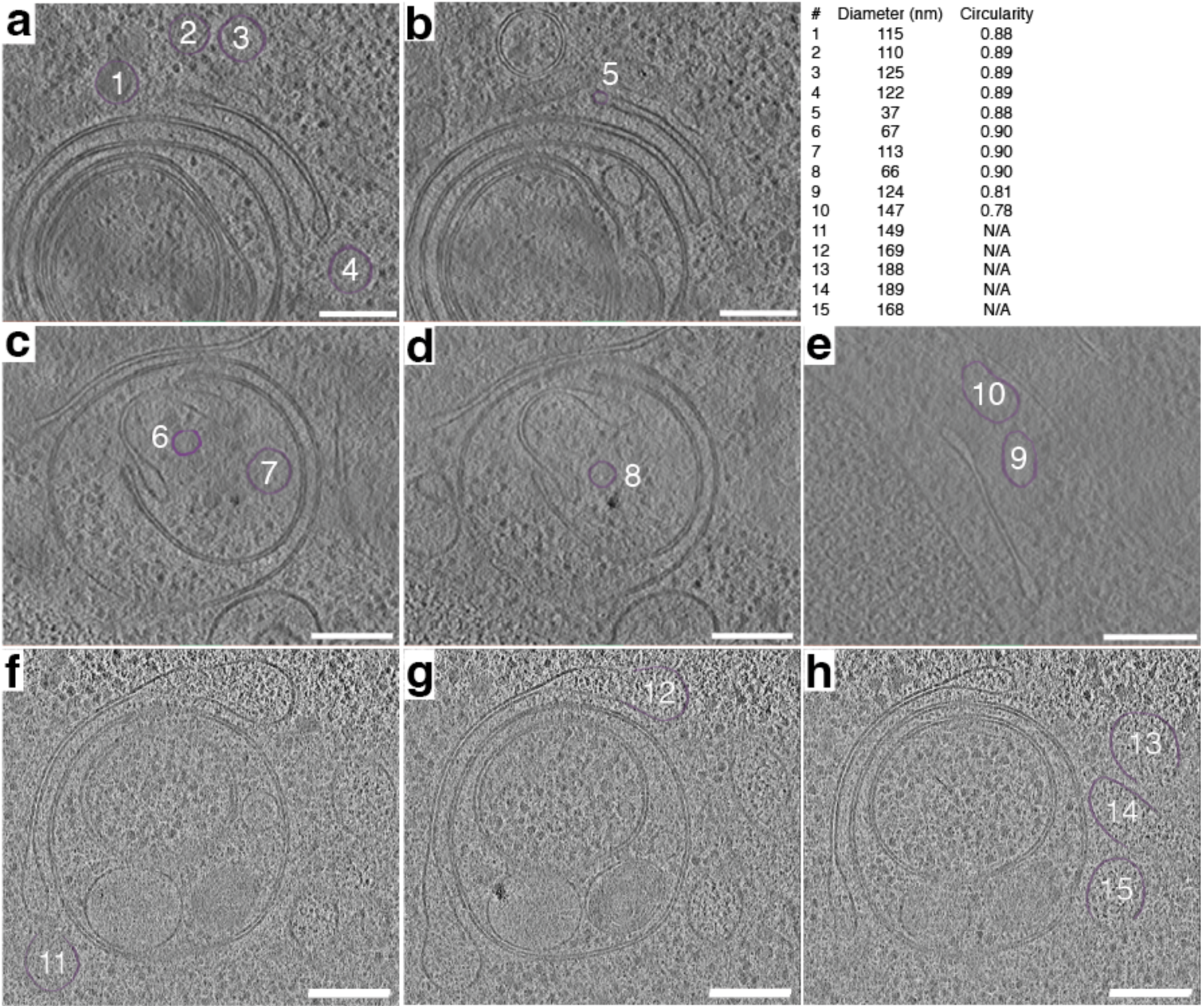
Tomographic slices of the Atg2-PM4 mutant cells. Vesicles proximal to phagophore rims are zoomed in slices of tomograms presented in Fig. 2, Fig.4 and Movie 1. Panels f)-g) show the fusion of vesicles with two phagophore rims. Circularity is defined here as 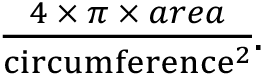 Diameters and circularity values can be found in Table 3. Scale bars 200 nm.

**Supplementary Figure 3:**
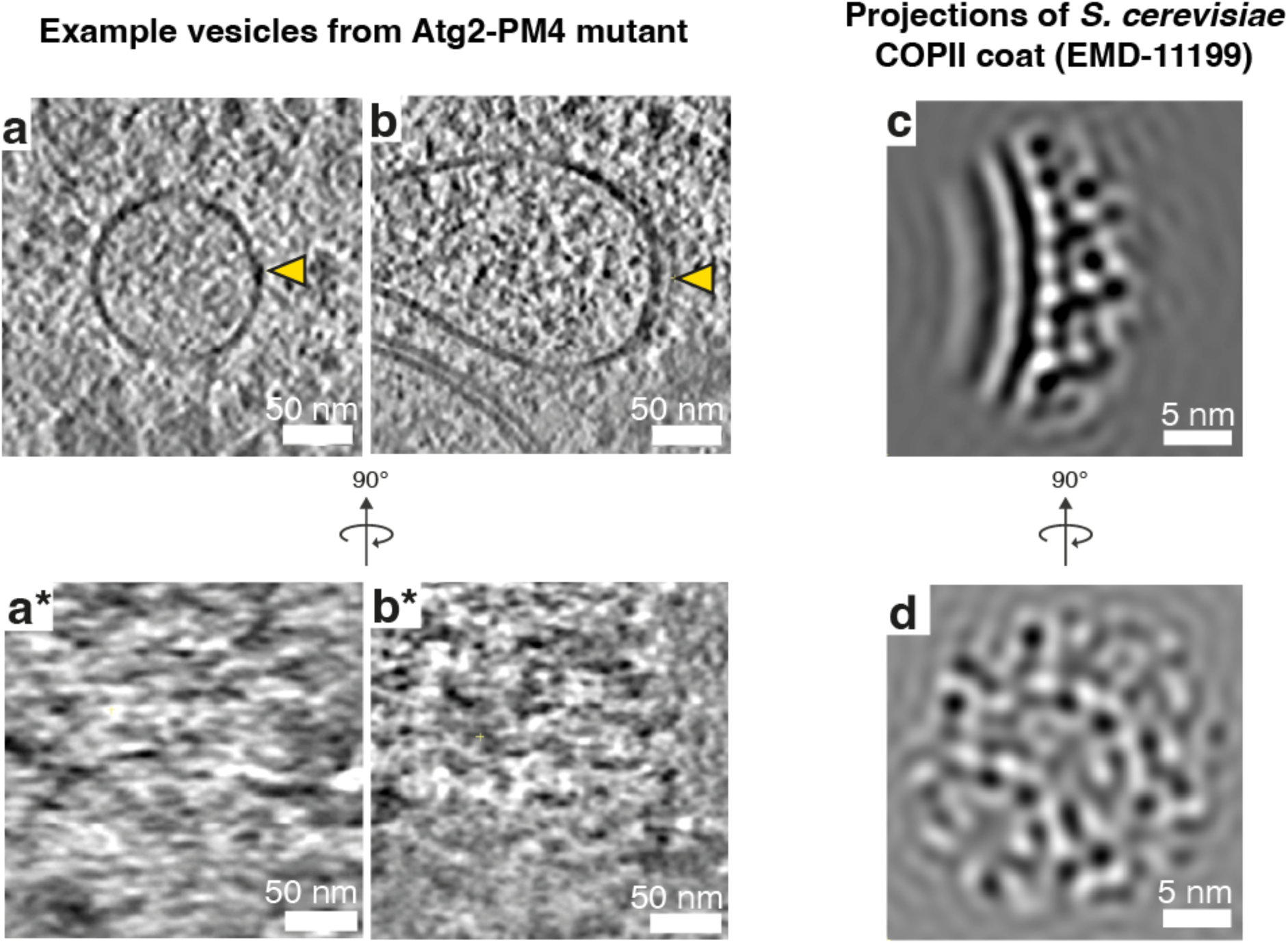
Comparison of detailed vesicle surface features with reference COPII vesicles. **a**) – **b**) Example vesicles from the Atg2-PM4 mutant. Yellow arrow heads indicate the side from which the cross section in panels a* and b* were taken. **a\***)-**b\***) Cross-section of the region above the vesicles where a protein coat lattice should be visualizable. **c**) – **d**) Subtomogram average map projection from *S. cerevisiae* COPII protein coat generated by (Hutchings et al., 2021), low pass filtered to 20 Å and contrast inverted. Map was obtained from the EMDB (Entry: EMD-11199).

**Supplementary Figure 4:**
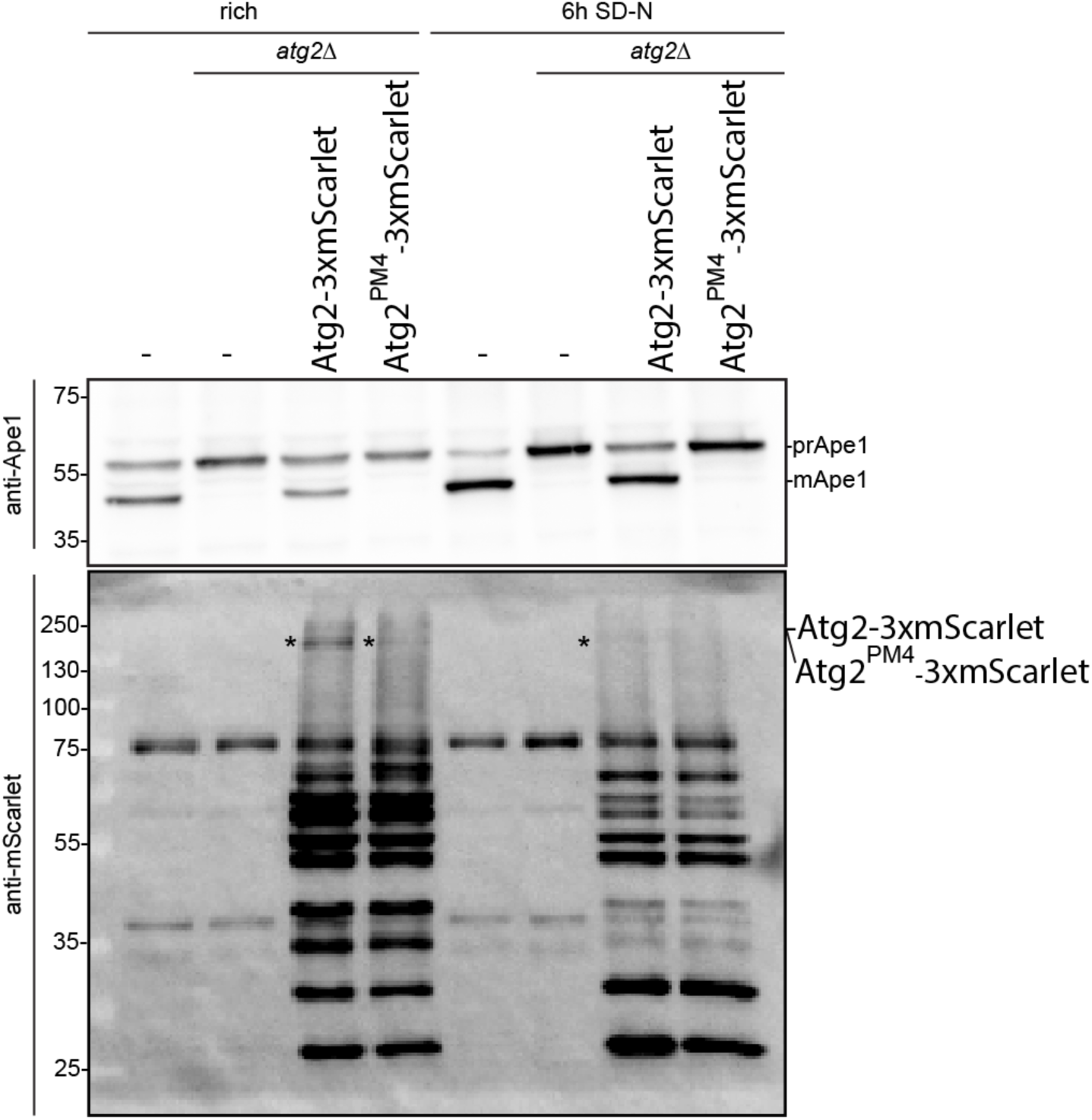
Characterization of Atg2-PM4 expression and autophagy flux. Western blot against Ape1 and Atg2 of lysate from wild type and *atg2Δ* (complemented with Atg2-3xmScarlet and Atg2-PM4-3xmScarlet) used in this study in synthetic media and under condition of 6 hours nitrogen starvation.

**Supplementary Table 1:**
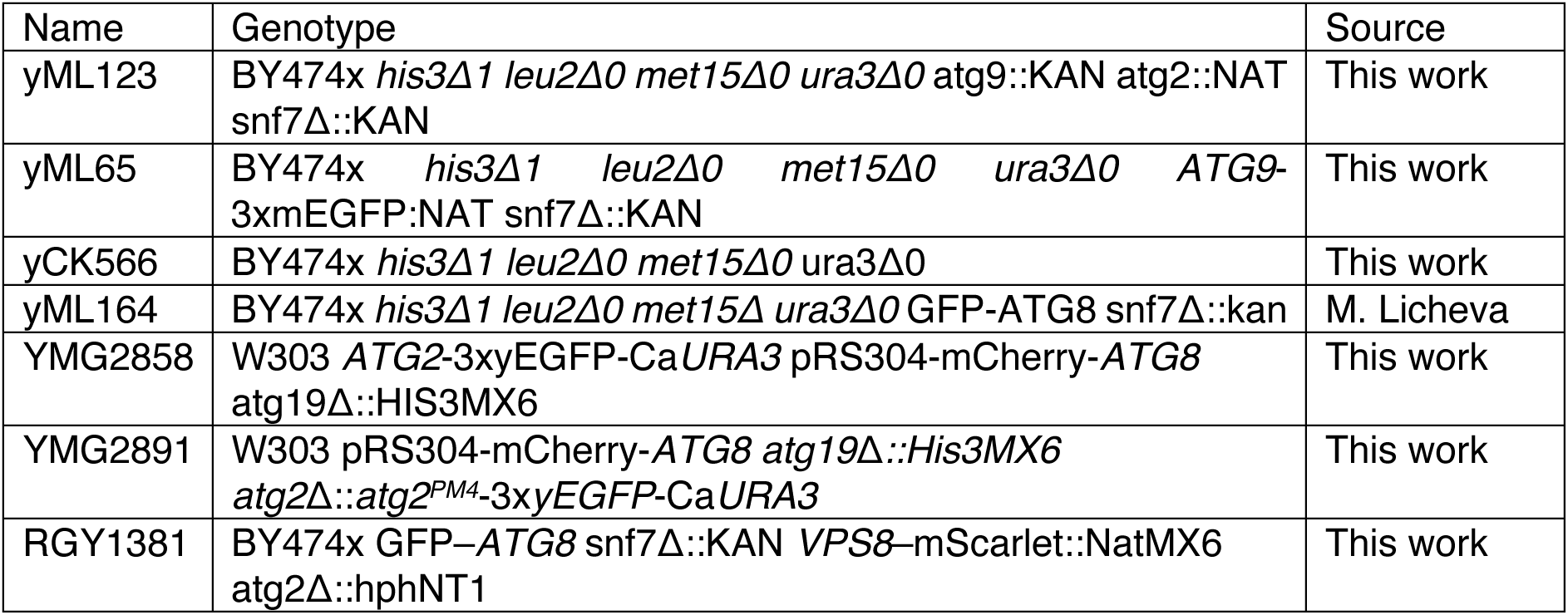
*S. cerevisiae* strains used in this study.

**Supplementary Table 2:** Plasmids used in this study.

| Name | Features | Plasmid backbone | Source |
| --- | --- | --- | --- |
| pCO114 | *atg2-promoter-atg2-3xmScarlet* | pRS315 | This work |
| pCO119 | *atg2-promoter-atg2<sup>PM4</sup>-3xmScarlet* | pRS315 | This work |
| pCO118 | *atg9-promoter-atg9-3xmTagBFP2* | pRS415 | This work |
| pRS416 | - | pRS416 | (Sikorski and Hieter, 1989) |
| pRGS277 | *atg2-promoter-atg2* | pRS416 | This work |
| pRGS278 | *atg2-promoter-atg2<sup>PM4</sup>* | pRS416 | This work |
| EMG841 | *atg2 promoter-atg2<sup>PM4</sup>-3xGFP-CaUraMX6* | pRS315 | This work |

